# Multiparametric Characterization of Individual Suspended Nanoparticles using Confocal Fluorescence and Interferometric Scattering Microscopy with Microfluidic Confinement

**DOI:** 10.1101/2025.07.25.666405

**Authors:** Eric Boateng, Erik Olsén, Albert Kamanzi, Yao Zhang, Bin Zhao, Pieter R. Cullis, Sabrina Leslie

## Abstract

Detailed characterization of the size, mass, payload, and structure of suspended mRNA-lipid nanoparticles (LNPs) is necessary to improve our understanding of how these heterogenous properties influence therapeutic efficacy and potency. Methods currently in use face limitations in reporting ensemble-average particle properties or requiring dedicated home-built microscopes that are beyond the reach of nanoparticle developers. In this work, we overcome these limitations by combining a commercially available confocal microscope and a convex lens-induced confinement (CLiC) instrument to achieve simultaneous characterization and correlation of the size, mass, refractive index, and nucleic acid payload of individual LNPs. We established the accuracy and precision of our method using nanosized beads and used it to investigate the size, payload, and water content of LNPs in different solvent pH. By employing readily available microscopy tools, we open the door to widespread adoption of our quantitative, in-solution nanoparticle characterization method.

---

Detailed information on the size, mass, and payload of heterogeneous nanoparticle suspensions is critical for the advancement of nanoscience, as well as for clinical and industrial applications[1, 2]. In recent decades, optical characterization techniques have been established as complementary high-throughput alternatives to electron microscopy by allowing detailed information about individual nanoparticles to be obtained under physiologically relevant conditions[3–7]. Among these, label-free methods such as evanescent scattering and interferometric scattering (iSCAT) microscopy have enabled quantitative measurements of individual nanoparticles and proteins[8–12]. Similarly, modern fluorescence-based microscopy techniques can detect and quantify the signal of individual fluorophores that label specific moieties in a mixture[6, 13]. When combined with single-particle tracking, these optical microscopy techniques can quantify the distribution of particle sizes through single-particle diffusivity measurements[14]. The relationship between size, intensity, and mass provides additional information on structure and material composition[5, 6, 15, 16]. The accuracy and precision of such measurements ultimately depend on measurement parameters and statistics, including the number of single-particle trajectories, track lengths, and the signal-to-noise ratio (SNR) for which each particle is detected, and other aspects of the image quality.

Although optical methods for nanoparticle characterization are increasingly being adopted, their widespread implementation has been limited by practical challenges that often require specialized imaging setups. One key challenge lies in accurately characterizing suspended nanoparticles in solution, especially in terms of both size and optical signal. Recent advances in label-free imaging, particularly the use of iSCAT, have addressed this limitation, enabling precise label-free characterization of biological nanoparticles[5, 9]. A second challenge is that most characterization methods have thus far involved home-built optical setups, which are often out of reach for nanoparticle developers. It has, however, recently been shown that a commercial confocal microscope can be used for fluorescence-based snapshot single-particle characterization of freely diffusing lipid nanoparticles (LNPs) containing mRNA in terms of mRNA loading and membrane fluidity[17, 18]. A third challenge is achieving simultaneous, complementary, and quantitative fluorescence and label-free light-scattering insights from suspended single-particle measurements over time – where fluorescent labels provide specificity for multi-valent cargo characterization, and the label-free signals allow for mass and size determination. Due to effects such as out-of-focus bleaching, quantitative single-nanoparticle imaging using both label-free and fluorescence microscopy has hitherto been restricted to surface-based methods[7, 19, 20]. However, tethering of nanoparticles to a surface is known to introduce biases owing to surface interactions, tethering chemistry, or steric constraints on conformations. Thus, overcoming all of these challenges to enable multiparametric single-particle characterization of suspended nanoparticles remains a significant need.

In this work, we address these single-nanoparticle imaging challenges by combining an inverted confocal microscope with a microfluidic Convex Lens-Induced Confinement (CLiC) instrument[6, 21]. We can perform simultaneous multicolor (MC) fluorescence and label-free iSCAT imaging of suspended nanoparticles without out-of-focus bleaching by trapping individual particles in microwells, herein referred to as CLiC-MC-iSCAT (Figure 1 A-B). The nanoparticle suspension is loaded into a CLiC flow cell containing an embedded array of microwells (Supporting Information, Section 1.5). Each microwell is 500 nm deep, matching the focal depth, and 3 µm in diameter, serving to isolate individual particles and enable extended single-particle trajectory measurements while keeping them in focus[6]. From these trajectories, single-particle sizes are obtained by fitting the resulting mean square displacement data to a 2D confinement model (Figure 1C). This enables size determination in addition to quantification of the polarizability and fluorescent cargo from the iSCAT and fluorescence images.

**Fig. 1.**
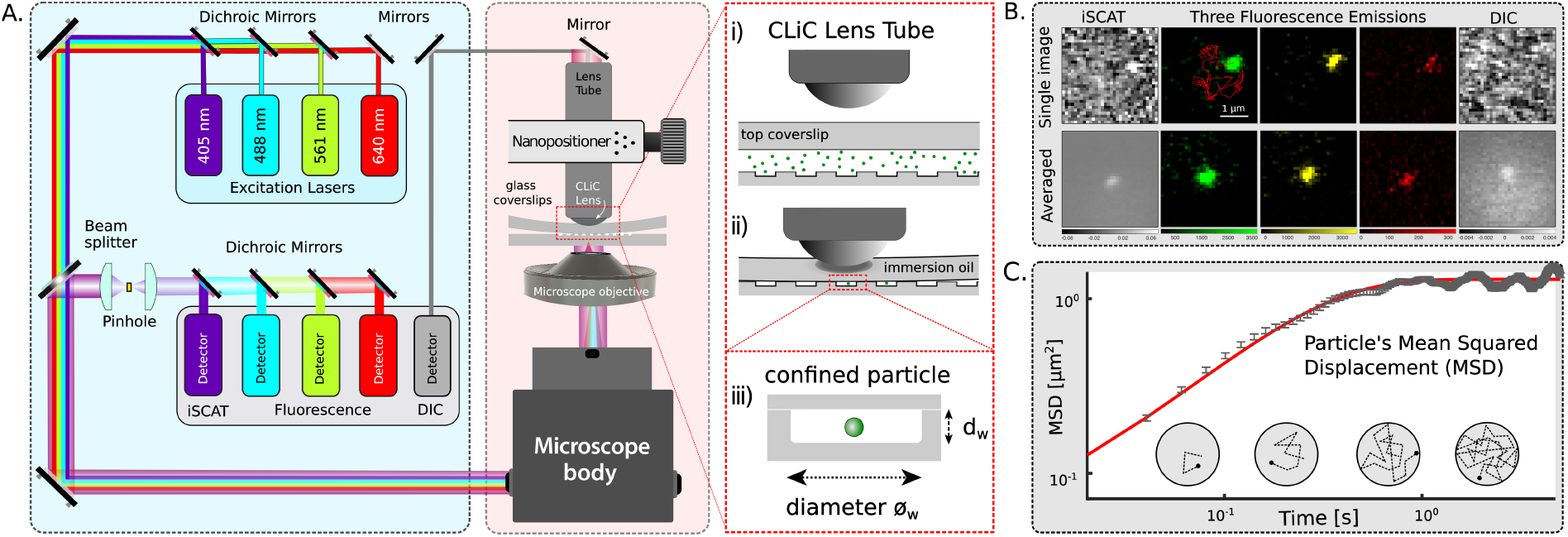
Overview of confocal CLiC-MC-iSCAT imaging. **A**. Schematic of the confocal microscope equipped with four excitation lasers and five detectors, where the CLiC instrument is placed on top of the microscope stage. The 405 nm laser is used for iSCAT imaging and the other three wavelengths (488, 561, 640 nm) are used for fluorescence imaging. Any of the lasers could be used for simultaneous differential interference contrast (DIC) transmission imaging. The CLiC lens tube is controlled by piezo actuators which push the CLiC-lens down onto the top coverslip, deforming it until it makes contact with the bottom coverslip containing an embedded array of microwells. Each microwell has a 3 µm width and 500 nm depth. The concentration is sufficiently dilute so that most wells contain either zero or one particle. After the imaging is performed, the top coverslip is raised which creates the opportunity to re-populate the wells and repeat the measurement process, thus increasing measurement statistics. **B**. Simultaneous fluorescence and label-free imaging where each particle is detected in at least one fluorescence channel in addition to iSCAT and DIC channels. Particle tracking is performed using one selected fluorescence channel, and for each particle detection, a centered region of interest image is extracted and saved from all channels for further image processing (shown here for a 100 nm polystyrene bead). The absolute values of the per-particle regions of interests are then averaged across detections to improve the image SNR during colocalization and signal quantification. **C**. The single-particle trajectories are converted to a mean-squared displacement curve from which the single-particle diffusivity and hydrodynamic diameter are quantified using confined diffusion theory (Supporting Information, Section 1.9)[6].

The imaging setup is a Nikon AXR confocal microscope equipped with 405, 488, 561, and 640 nm lasers. Simultaneous fluorescence and label-free imaging was achieved with minor modifications to the microscope (Supporting Information, Section 1.6). The 405 nm laser was used for iSCAT imaging, combinations of the other three lasers were used to detect fluorescence markers, and any of the lasers could be used for differential interference contrast (DIC) imaging in transmission mode (Figure 1A). The simultaneously measured iSCAT signal amplitude and size obtained from particle tracking analysis were combined to estimate the particle refractive index using Mie theory (Supporting Information, Section 1.8). The refractive index provides information on the particle material[5, 15], which in the context of LNPs can be related to the amount of water within the particles[22]. To minimize potential surface interactions, the flow cells were prepared either using piranha-glass cleaning protocols for reference bead samples or surface PEGylation protocols for LNP formulations developed in prior work[6] (Supporting Information, Section 1.5).

An advantage of using CLiC for simultaneous multichannel imaging of a single particle is that the particle remains in focus at all times, allowing particle tracking to be performed in only one of the imaging channels. The position of the particle in each frame is used to crop the corresponding images in the other channels, where the crops are used to evaluate the particle signal and potential colocalization. The array of cropped images, which are centered on the same single-particle trajectory, can then also be processed (e.g. averaged, Figure 1B) to improve the SNR during subsequent quantification steps. Averaging across frames is particularly beneficial for label-free images or slowly bleaching fluorophores, as all or most particle observations can then be used during the averaging. For multicolor fluorescence, only fluorescence signals that move together with the LNP during the imaging will create a clear spot in the averaged image, whereas signals that do not move together will only create a background signal in the averaged image (Supporting Information, Section 1.7.3). For the label-free images, the absolute contrast is used during the averaging to account for the depth-dependent variations in relative phase difference between the particle and the background (Supporting Information, Section 1.7.2)[23]. For all of the particle measurements in this work, one of the fluorescence channels was used for the single-particle tracking. The signals of each particle were quantified by applying a Gaussian fit to an averaged image of a series of cropped images centered on particle positions along its trajectory (Supporting Information, Section 1.7). In a representative experiment, when imaging 100 nm diameter polystyrene nanoparticles labeled with multiple fluorophores, each particle was tracked and its reference position was determined using only a single fluorescence channel. 100% colocalization was achieved with respect to the other imaging channels (Supporting Information, Figure S7).

As a first step to validate CLiC-MC-iSCAT, we measured several different fluorescently-labeled dielectric nanoparticles with known sizes and material properties. Figure 2A illustrates this simultaneous imaging in both fluorescence and label-free modalities. The observed scaling relationship between the iSCAT signal and polystyrene particles of different sizes was consistent with theoretical predictions. Moreover, the mean and standard deviation values obtained from the measured hydrodynamic diameters (49 ± 16, 82 ± 18, 119 ± 31, 244 ± 40 nm) were all similar to the size distributions obtained from complementary darkfield nanoparticle tracking analysis (NTA) measurements (51 ± 16, 78 ± 18, 122 ± 25, 260 ± 63 nm), described in the Supporting Information, Figure S2. These results indicate that the scanning speed of the confocal imaging system is sufficient for accurate nanoparticle size measurements using the mean square displacement curve and the theoretical model for confined particle diffusion in a well[6]. Moreover, the similarity in both the mean and width of the size distributions compared to NTA measurements suggests that the observed distribution widths are not due to potential surface interactions within the wells, but instead originates from a combination of finite track lengths and intrinsic sample heterogeneity (Supporting Information, Section 1.9).

**Fig. 2.**
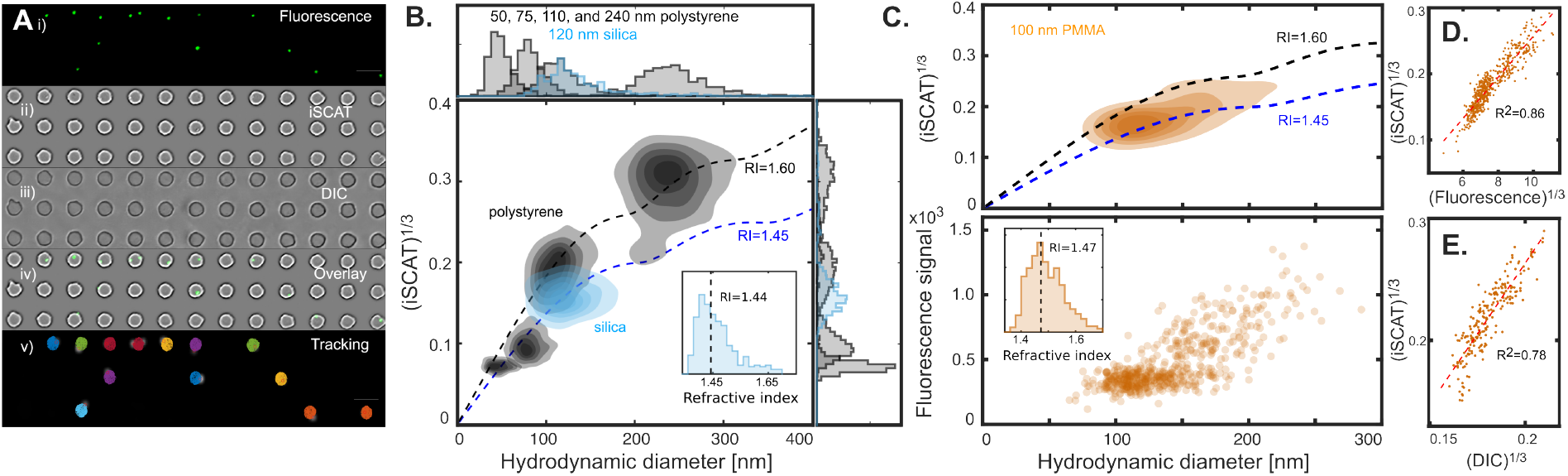
Quantitative validation of CLiC-MC-iSCAT imaging using dielectric nanoparticles. **A**. Illustration of the simultaneous CLiC-MC-iSCAT imaging of nanoparticles in both fluorescence and label-free. **i)**. Fluorescence emission of 100 nm polystyrene beads. **ii)**. iSCAT image of single bead particles confined in 3 µm microwells. Gaussian convolution of the images was performed to highlight the edges of the wells (Section 1.71). **iii)**. DIC image in forward transmission mode. **iv)**. Overlay image using both fluorescence and iSCAT channels showing colocalized particles confined in the wells. **v)**. Nanoparticle traces of individual confined particle to estimate single particle size via the diffusivity-size relation. Scale bar is 5 µm. **B**. The measured scaling relationship between the hydrodynamic diameter and the cube root of the iSCAT signal for a suite of nanosized polystyrene beads agrees well with predictions from Mie theory (black dashed line), which describes the scattering of light by small spherical particles (Supporting Information, Section 1.8). Different iSCAT signals were obtained for polystyrene and silica particles of the same size, which is consistent with the refractive index difference between silica and polystyrene. The inset is the estimated refractive index histogram for silica particles, where the estimated median refractive index is 1.44 (vertical dashed-black line from the inset image). The contour levels in the plot correspond to 17, 33, 50, 67, and 83% of the obtained distribution density. Each sample was measured separately. The number of particles in each distribution that is colocalized with iSCAT ranges from 300 particles for the 50 nm polystyrene to 1100 particles for the 100 nm polystyrene. **C-E** We demonstrate multi-parametric characterization of 100 nm diameter PMMA particles in a single measurement including: **C**. fluorescence and the cube root of the iSCAT scattering intensities as a function of hydrodynamic diameter, where both signals increase as a function of size. The black and blue dashed lines are Mie calculations for the refractive indices 1.60 and 1.45, respectively, to enable comparison with the results in **B**. Refractive index distribution of the measured particles, where the median refractive index is 1.47 (black dashed line). **D**. The cube root of the iSCAT signal plotted as a function of the cube root of the fluorescence signal, revealing a proportional relationship. This correlation is expected, as both signals, to a first approximation, scale with particle volume. **E**. The cube root of both the iSCAT signal plotted as a function of the cube root of the DIC signal, demonstrating a proportional relationship when both signals are detectable in respective imaging channels. Notably, for particles with lowest iSCAT signal, the corresponding DIC signal falls below detection and cannot be estimated.

To further study the relationship between size and iSCAT signal, we performed CLiC-MC-iSCAT measurements on nanosized particles of different materials, including polystyrene, silica, and polymethyl methacrylate (PMMA), where the estimated refractive indices all are close to the reference values (Figure 2). By using the 100 nm polystyrene measurement as a calibration point, we established that the scaling relationship between intensity and size for polystyrene particles is similar to what we expect from Mie theory (Supporting Information, Section 1.8). Moreover, the 100 nm diameter silica bead sample has a measured mean and median refractive indices of 1.47 ± 0.01 and 1.45 ± 0.01, respectively (Figure 2B), where the uncertainty here is the standard deviation of the mean. These estimates are close to previously reported value of 1.447 when using an illumination wavelength of 405 nm[24]. Similarly, our CLiC-MC-iSCAT characterization of 100 nm diameter PMMA nanospheres yields a mean and median refractive index of 1.49 ± 0.01 and 1.47 ± 0.01 (Figure 2C), which is in good agreement with the previously reported refractive index of 1.516 for PMMA[25].

In addition, we investigate how the iSCAT, DIC, and fluorescence signals scale with one another, as shown in Figure 2D-E for the PMMA particles. The observed correlations are consistent with expectations, as both DIC and fluorescence signals are known to increase with particle volume[6, 26]. This indicates that our observed spread in measured single-particle signals from a sample reflects the heterogeneity of the sample properties rather than the statistical uncertainty of the measurements.

Note that because the sensitivity of the fluorescence measurement is higher than that of the iSCAT measurement, as the particle size decreases, there is an increasing population of particles which can be reliably detected and tracked using fluorescence but not using iSCAT. For example, the measured polystyrene particles with diameter of 100 nm can be detected and tracked directly using the iSCAT signal on this microscope. For polystyrene with diameters of 40 nm or smaller, the iSCAT signal is below the detection limit, restricting the characterization of the hydrodynamic diameter of the particles and their fluorescence signals (Supporting Information, Figure S6). In between these sizes, the iSCAT signal can still be quantified because of the position-based averaging using the fluorescence signal. We note that a similar lower size limit in the detected concentration of the nanoparticles can be observed for the complementary darkfield and fluorescence NTA measurements (Supporting Information, Figure S2), highlighting that similar limitations exist for other widely used nanoparticle characterization methods.

Following these validation measurements, we used CLiC-MC-iSCAT to investigate the properties of Cy5-mRNA loaded LNPs with the green fluorescent lipophilic dye DiO as a lipid marker, where signal colocalization was used to evaluate the subpopulation of LNPs containing mRNA cargo[27, 28]. LNPs were formulated using the T-junction method, with a lipid composition of MC3, DSPC, cholesterol, PEG-DMG, and DiO at a molar ratio of 50/10/38.5/1.5/1.0, respectively, with a N/P ratio of 6 for the mRNA (Supporting Information Section 1.2). While LNP systems containing ionizable lipids are capable of encapsulating nucleic acids[29], not all LNPs within a sample necessarily contain mRNA cargo[28]. By tracking the LNPs using the DiO fluorescence signal, it is possible to determine the fraction of LNPs that contain Cy5-mRNA based on the evaluation of colocalization with the red Cy5 fluorescence emission at the single particle level. Further, the refractive index information from the iSCAT channel can be related to the water content within the LNP[22, 30], which is a common parameter used to understand the LNP structure[31].

Figure 3A shows single LNP measurements of mRNA loading distributions as a function of particle size, characterized by the ratio of Cy5 to DiO signals measured per particle. Our measurements show two distinct clusters of loaded versus unloaded LNPs, with approximately 73.7% of the particles having Cy5-mRNA signal after correcting for potential channel cross-talk (Supporting Information, Section 1.7). The diameter of the LNPs containing mRNA is 99 ± 46 nm (mean and standard deviation), which is on average 19 nm larger than particles without mRNA (79 ± 32 nm). Notably, our observation that ∼ 25% of LNPs do not contain mRNA cargo is within the range of previously reported loading fraction values for MC3-based LNPs[17, 28]. Moreover, the observed increase in particle size for LNPs containing mRNA cargo is consistent with prior work that has shown that LNPs made without the presence of mRNA are typically smaller than those formulated with the presence of RNA[32].

**Fig. 3.**
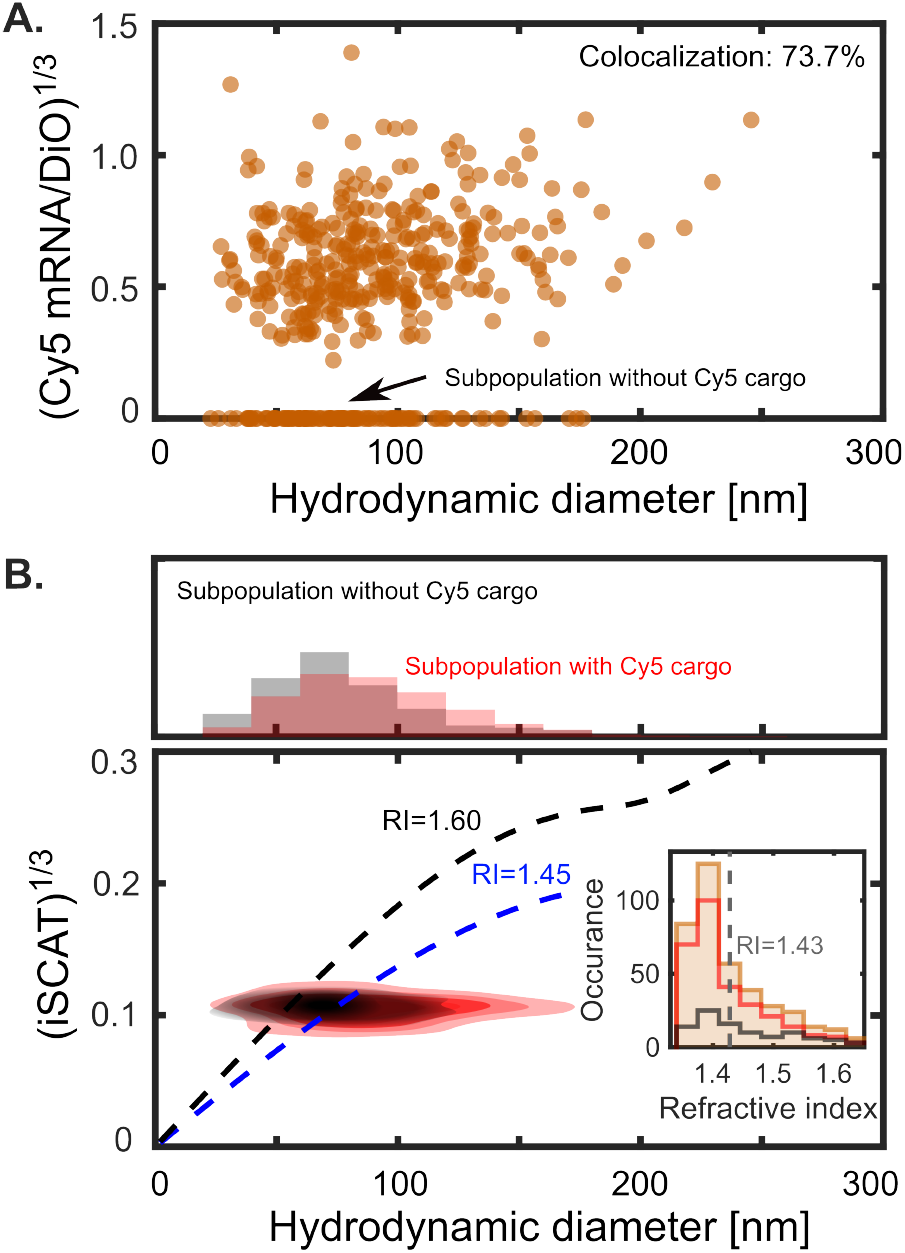
CLiC-confocal iSCAT and MC fluorescence characterization of mRNA lipid nanoparticles. CLiC-MC-iSCAT measurements of LNPs containing MC3, DSPC, cholesterol, PEG-DMG, and DiO, at a molar ratio of 50/10/38.5/1.5/1.0, respectively, with a N/P ratio of 6 for the mRNA. **A**. Scatter-plot of the cube root of the Cy5 mRNA-DiO lipid signal, used to identify particles with and without Cy5 cargo. **B**. Size-cube root of the iSCAT signal contour plot of the mRNA-LNPs, where the LNPs colocalized with Cy5-mRNA cargo are in red, whereas those without Cy5-mRNA signal are in black. The contour levels correspond to 17, 33, 50, 67, and 83% of the obtained distribution density. The inset is the refractive index histogram of the LNPs, where the median refractive index of 1.43 indicates a water content around 40% inside the LNPs (gray dashed line). The solid red and black lines in the inset correspond to the refractive index distribution of LNPs colocalized with and without mRNA.

Following the analysis of the fluorescence signals, we combine the estimated particle size and iSCAT signal to determine that the median refractive index of the measured LNPs is 1.43 ± 0.01 (Figure 3B). This agrees well with previously reported estimates of the LNP refractive index[22, 30]. A typical mRNA-LNP is made up of three main components: water (refractive index of 1.34), lipids (refractive index = 1.49-1.50[33]) and mRNA (refractive index = 1.61-1.63[34]), where we estimate these values using an imaging wavelength of 405 nm. By assuming that about 10% of the biomolecules in the LNP correspond to mRNA and the rest correspond to lipid[31] molecules, the average biomolecular refractive index *n*_bio_ can be − approximated as ∼1.51-1.52. For biological particles, their refractive index *n*_p_ generally scales linearly with biomolecular concentration[35], *n*_p_ − *n*_media_ = (*n*_bio_–*n*_media_)*V f*_bio_ where *V f*_bio_ is the volume fraction of the biomolecules. Following this reasoning, the average refractive index can be related to an average LNP water content of ∼50%. Note that this water content refers to the full LNP, including the LNP, outer PEG layer, and the hydration layer, whereas other water fraction estimates typically only refer to the core of the LNP[31]. The inclusion of the outer PEG layer and the hydration layer is the reason for the higher reported value compared to the values of the water fraction only referring to the core of the LNPs (Supporting Information, Section 1.10). It can also be noted that the particles colocalized with mRNA have a similar iSCAT signal as the particles without colocalized mRNA while having a larger particle size. This in turn implies that LNPs colocalized with mRNA have a lower refractive index than those without colocalized mRNA, which indicates a higher water content for those LNPs, particularly considering that mRNA has a higher refractive index than lipids. Given the presence of blebs in the cryoTEM data (Supporting Figure S1), where blebs are associated with a water phase inside LNPs where the mRNA is residing[28], it is likely that the water inside the bleb counteracts the higher refractive index of the mRNA (Supporting Information, Section 1.10). However, validation data would be needed to evaluate this observation.

In addition to particle size and mRNA loading, another important property of LNPs is their response to a reduced surrounding pH[36, 37]. To further evaluate CLiC-MC-iSCAT and the ability to quantify multiple signals on a single-particle level, we investigated LNPs containing a cargo of DNA with a conjugated pair of pH-sensitive and pH-insensitive dyes[37]. The two fluorescence signals, when measured and analyzed simultaneously on the single particle level, can be used to monitor intraparticle pH in real time (Supporting Information, Section 1.3)[37]. The pair of dyes consists of Alexa647 and FAM (6-carboxyfluorescein), where the emission of Alexa647 is pH insensitive, while the fluorescence emission of FAM decreases as the pH is lowers[37].

When comparing LNP measurements initially performed at neutral pH (1× PBS buffer, pH 7.4) and then repeated under acidic conditions (sodium acetate buffer, pH 4) (Supporting Information, Section 1.7), we observed that when lowering the pH of solution the FAM/Alexa647 ratio decreases while the size of the individual LNPs on average increase (Figure 4). The reduction in the fluorescence signal ratio of the particles is expected, highlighting that the fluorescence signal estimation can be compared accurately between different sample measurements, where a signal reduction by a factor of 4 has previously been measured at the ensemble level[37]. The estimated median particle refractive index at neutral pH is around 1.42 ± 0.01, which is similar to that of the mRNA LNPs (Figure 3C). When lowering the surrounding pH, the increase in LNP size reduces the estimated refractive index to around 1.38 ± 0.01, which corresponds to an increase in the water content inside the LNPs from around 50% to around 75%. Since a similar size increase is observed 7 using DLS (Supporting Information, Figure S8), the increased estimated particle size is not due to potential pH-dependent surface interactions when the particles are in the microwells. Moreover, since the measured iSCAT signal remains similar in the two different pH environments, this indicates that the increase in size is not due to particle aggregation as that would have caused the measured iSCAT signal to increase with increasing particle size. Instead, the combination of a slight reduction in the scattering signal with an increase in size upon lowering the pH is consistent with LNP swelling (Supporting Information, Section 2.3). This observation agrees with the protonation behavior of the ionizable lipid in previous studies[38], in which it has been hypothesized that electrostatic repulsion between charged MC3 lipids at reduced pH triggers a structural change in which the resulting influx of water and ions balances the repulsion.

**Fig. 4.**
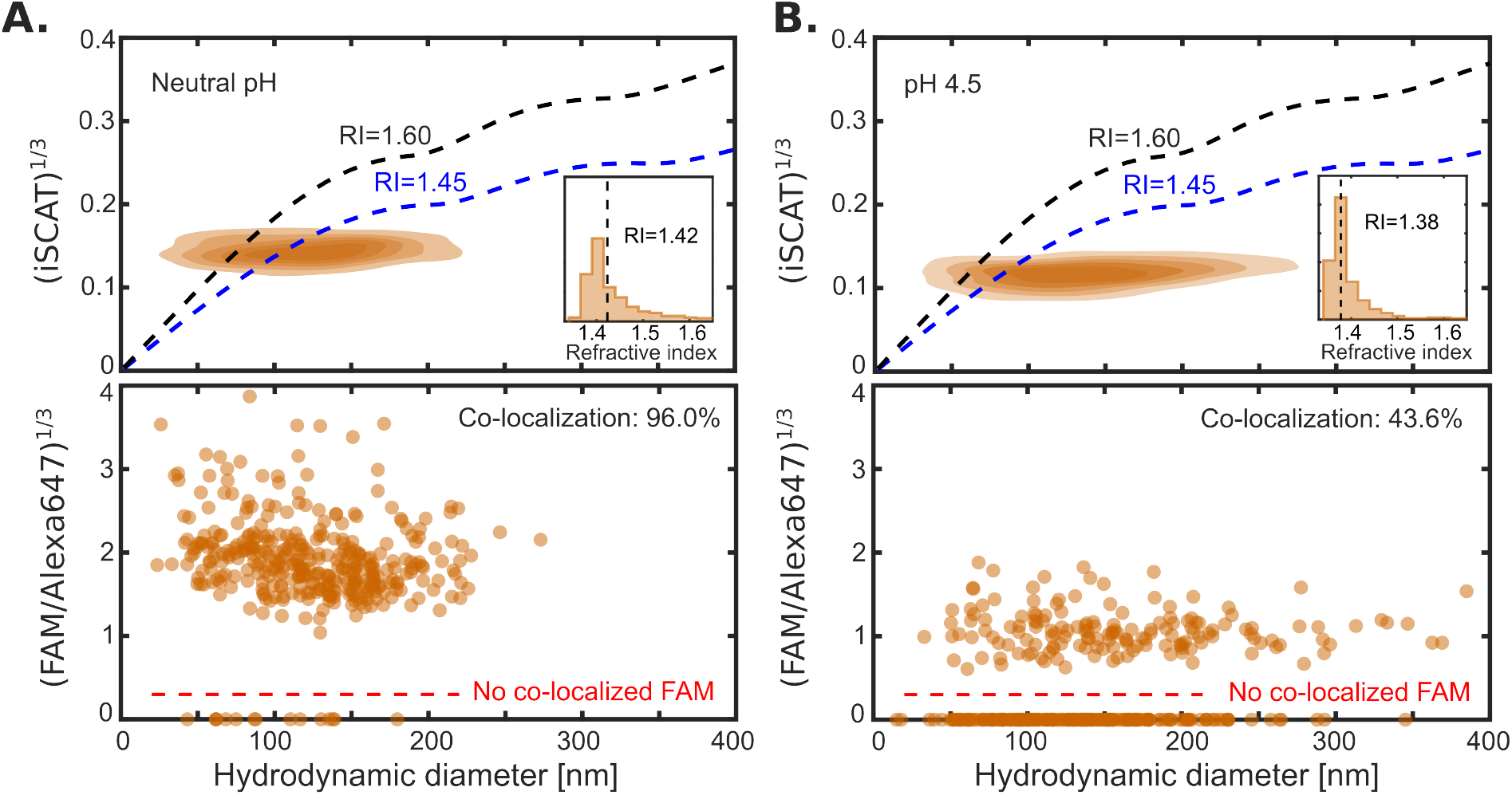
Multi-parametric characterization of suspended LNPs at different pH, containing DNA cargo which is labeled with both pH-sensitive and pH-insensitive dyes. CLiC-MC-iSCAT measurements of LNPs containing MC3, DSPC, cholesterol, and PEG-DMG at a molar ratio of 50/10/39/1.0, respectively, with a N/P ratio of 3 for the DNA cargo. **A**. Measurements of LNPs at neutral pH in a PBS buffer. The cube root of both the iSCAT signal and fluorescence FAM/Alexa647 ratio as function of size, where almost all detected LNPs (96 %) had a co-localized FAM signal. The contour levels correspond to 17, 33, 50, 67, and 83% of the obtained distribution density. The inset is a histogram of the single-particle refractive index, where the median refractive index of the LNPs is 1.42 (black dashed line), which is similar to that of the mRNA LNPs. **B**. Measurements of LNPs at reduced pH in an acetate buffer. The cube root of both the iSCAT signal and fluorescence FAM/Alexa647 ratio as function of size. We note that the estimated hydrodynamic diameter is noticeably larger than that for neutral pH, and the fluorescence ratio is reduced such that only 43.6% of all LNPs are co-localized with a FAM signal. The inset is a histogram of the single particle refractive index, where the median refractive index of the LNPs is 1.38 (black dashed line).

In conclusion, we have introduced and established CLiC-MC-iSCAT as a versatile method for multiparametric particle characterization. Specifically, CLiC-MC-iSCAT enables the measurement of size, loading, and refractive index of suspended nanoparticles using a multicolor confocal microscope combined with the CLiC instrument. We have benchmarked our size and iSCAT signal measurement approach using standard nanosized beads and established agreement with predictions from Mie theory. This has enabled us to combine and interpret our measurements of the iSCAT signal and to estimate the refractive index of each particle in a heterogeneous suspension as all particle estimates are here made on the single-particle level. Furthermore, we have applied our simultaneous measurements of multichannel fluorescence and iSCAT, in order to study the change in LNP properties as a function of pH. The information obtained from CLiC-MC-iSCAT measurements on individual particles is extensive and complementary to what is possible to measure using most standard characterization techniques (Supporting Information, Section 1.11). Furthermore, given the usage of confocal microscopes to investigate cells[39, 40], CLiC-MC-iSCAT can be used to obtain quantitative reference data of particles in solution on the same microscope as in live cell experiments. Thus, we anticipate that this type of optical-microscopy-based multiparametric characterization will find widespread applications and have a high impact in advancing many areas where suspended nanoparticles play an important role, ranging from industrial processes to drug discovery, cell measurements, and medical diagnostics.

## Supporting Information

Materials and methods, details regarding data analysis and how the particle measurements are related to particle properties, and supplementary figures, including cryo-TEM of LNPs (Figure S1), complementary NTA data (Figure S2), additional methodology details (Figures S3-S6), and measurement of reference beads containing fluorophores of multiple colors (Figure S7).

Supplementary Movie 1

Supplementary Movie 2

Supplementary Movie 3

Supplementary Movie 4

Supplementary Movie 5

## Acknowledgments

We acknowledge the substantial financial and other support from several granting agencies, including the Canadian Foundation for Innovation, British Columbia Knowledge Development Fund, PacifiCan and Western Diversification Fund, National Science and Engineering Research Council of Canada, Nanomedicines Innovation Network, as well as Startup Funds from the University of British Columbia Faculty of Science, Department of Physics and Astronomy, and Michael Smith Labs. Moreover, the Mitacs granting agency and industry sponsor ScopeSys co-supported two fellowships: a Graduate fellowship held by E.B. and Postdoctoral fellowship held by A.K.. Further, E.O. held an international Postdoctoral fellowship from the Swedish Research Council (grant number 2024-00439) and Y.Z. held a NanoMedicines Innovation Network graduate award and a Canadian Institutes of Health Research Doctoral Award (FBD 193487). S.L. was supported by a Killam Accelerator Research Fellowship. We extend our gratitude to the Nikon Canada staff who assisted with the setup of the microscope. We are thankful for the contributions from several undergraduate summer students, technicians, and operational and administrative staff throughout this work. We thank Dr. Edward Grant for supporting the early development and iSCAT training of E.B. Cryo-TEM data and grids were prepared at the High Resolution Macromolecular Electron Microscopy facility at the University of British Columbia, where we thank the staff and faculty of this facility.

## Notes

The authors declare the following competing financial interest(s): A.K. and S.L. have interest in ScopeSys Inc., a UBC spin-off company that is commercializing the CLiC technology. P.R.C. has interest in several companies that are commercializing mRNA LNP technologies, including Nanovation, Acuitas and more than 10 others.

