## Supporting Information for "Multiparametric Characterization of Individual Suspended Nanoparticles using Confocal Fluorescence and Interferometric Scattering Microscopy with Microfluidic Confinement"

**2 Complementary data** **19**
2.1 CLiC-MC-iSCAT size distribution of the measured reference beads . . 19
2.2 Evaluation of CLiC-MC-iSCAT using multi-labeled reference beads . . 20
2.3 Effect on measured size and iSCAT signal from potential particle swelling 20

### 1 Materials and methods

#### 1.1 Particle materials

Fluorescent reference dielectric nanoparticles; polystyrene beads of varying sizes (0.03±0.0034, 0.04±0.0043, 0.05±0.0160, 0.08±0.0038, 0.11±0.0089, 0.22±0.0079 µm diameter 505/515 nm ex/em FluoSpheres; F8787, F8795, F10720, G75, F8803, F8811) were purchased from ThermoFischer, Green silica beads (100 nm diameter, 488/518 nm ex/em, ASFG-21-0010) and Green PMMA fluorescent particles (100 nm diameter 488/518 nm ex/em, APFG-21-0010) were purchased from Abvigen Inc. The 100 nm diameter TetraSpeck beads (0.11 ± 0.009) used to evaluate the channel colocalization were purchased from ThermoFischer.

All lipid nanoparticles (LNPs) in the main text contain MC3/DSPC/cholesterol/PEG-DMG, where the mRNA-containing LNPs also contain DiO as a lipid dye. The ionizable lipid DLin-MC3-DMA (MC3) was purchased and synthesized by Dr. Marco Ciufolini. The lipids DSPC and PEG-DMG were purchased from Avanti Polar Lipids. Cholesterol was purchased from Sigma-Aldrich. The lipophilic dye, DiO, was purchased from ThermoFisher Scientific. For the LNP cargo, firefly Luciferase Cy5-mRNA was purchased from ApexBio, and all DNA oligonucleotides were synthesized and purified by Integrated DNA Technologies.

#### 1.2 Formulation of LNPs containing mRNA

Lipid components consisting of MC3/DSPC/cholesterol/PEG-DMG/DiO at a molar ratio of 50/10/38.5/1.5/1, respectively, were dissolved in the ethanol phase at a concentration of 10 mM total lipids. The lipid mixture was then rapidly mixed with an aqueous solution of Cy5-mRNA in pH 4 buffer (300 mM sodium citrate). The amine-to-phosphate ratio (N/P) was maintained at 6 for all mRNA LNP formulations. Rapid mixing was performed with a T-junction mixer at a 1:3 ratio of ethanol/aqueous buffer (v/v) and a final flow rate of 20 ml/min. The resulting dispersion was then dialyzed overnight against at least 500-fold volume of pH 7.4 PBS buffer. The prepared LNPs were then sterile filtered with a 0.2 μm Supor membrane syringe filter (Pall Corporation, Mississauga, ON, Canada).

#### 1.3 Formulation of LNPs for pH measurements

A pair of pH-sensitive and pH-insensitive dyes, 6-FAM and Alexa Flour 647, were chemically conjugated in a DNA duplex to form a 6-FAM&Alexa 647-DNA complex. This 6-FAM&Alexa 647-DNA complex is composed of O1-6FAM (6-FAM - TACATTTTACGCCTGGTGCCT), O2-A647 (CCGACCGCAGGATCCTATAA - Alexa Fluor 647), and O3 (TTATAGGATCCTGCGGTCGGAGGCACCAGGCG-TAAAATGTA), which was prepared as previously described[1].

The LNPs containing 6-FAM&Alexa Flour 647-DNA complex were formulated by first dissolving the lipids (MC3, DSPC, cholesterol, and PEG-DMG) into ethanol to a final concentration of 10 mM and at a ratio of 50/10/39/1 mol, respectively. The lipid mixture was then rapidly mixed with an aqueous solution containing 6-FAM&Alexa647-DNA complex in NaOAc-NaCl (25 mM sodium acetate, 150 mM sodium chloride) pH 4 buffer using a T-junction mixer at a 1:3 ethanol/aqueous buffer ratio (v/v) and a final flow rate of 20 ml/min. The amine-to-phosphate ratio (N/P) was 3. The resulting mixture was dialyzed overnight against at least 500-fold volume of pH 7.4 PBS buffer. The prepared LNPs were then sterile filtered with a 0.2 μm Supor membrane syringe filter (Pall Corporation, Mississauga, ON, Canada).

#### 1.4 Complementary particle analysis

##### 1.4.1 Sample analysis of lipid nanoparticles

Lipid concentrations were determined using the Cholesterol E Kit (Wako Diagnostics). Bulk particle size measurements were performed through dynamic light scattering using a Zetasizer Nano ZS (Malvern Instruments Inc.).

The mRNA-containing LNPs were also imaged using cryogenic transmission electron microscopy (cryo-TEM). LNPs were concentrated to 15 - 20 mg/ml total lipids using 30K Amicon ultracentrifugation units (EMD Millipore Corporation, Billerica, MA, USA). The particles were then added (3 - 5 µl) to glow-discharged copper grids and plunge-frozen with an FEI Mark IV Vitrobot. All imaging was performed with a FEI LaB6 G2 TEM operating at 200 kV low-dose conditions with an FEI Eagle 4K CCD camera. The samples were visualized at 55,000 times magnification with a nominal under focus of 1-2 µm to enhance contrast. Cryo-TEM imaging was performed by the High Resolution Macromolecular Cryo-Electron Microscopy Facility at the University of British Columbia (UBC). An example of the MC3-mRNA LNPs can be seen in Figure S1. Note that all LNPs are not necessarily spherical as some LNPs can contain blebs[2].

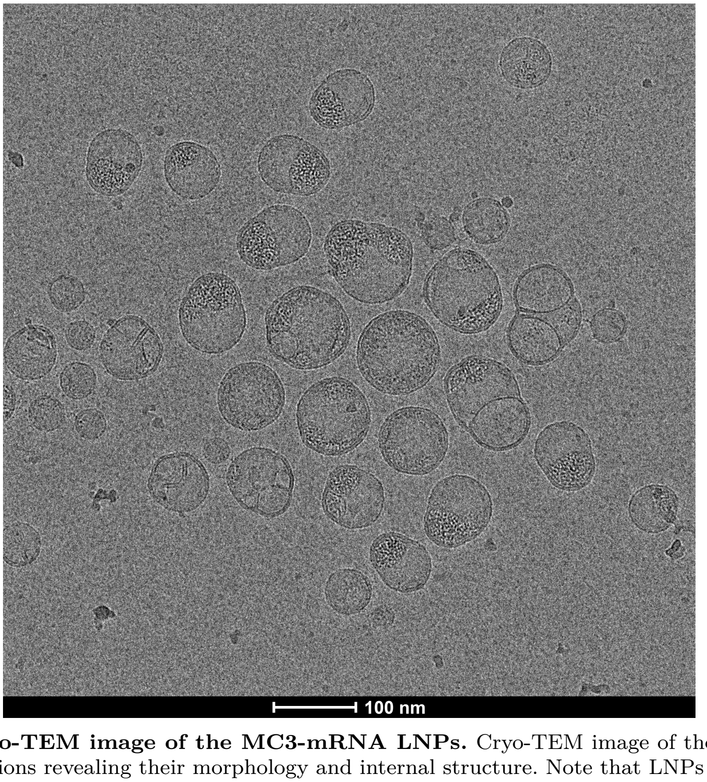

**Fig. S1 Cryo-TEM image of the MC3-mRNA LNPs.** Cryo-TEM image of the MC3-mRNA LNP formulations revealing their morphology and internal structure. Note that LNPs are not necessarily spherical as they can contain blebs[2].

##### 1.4.2 Sample analysis of reference particles using NTA

Complementary size distribution measurements of the reference particles were performed using Nanosight Pro from Malvern, which is a nanoparticle tracking analysis (NTA) instrument that measures the hydrodynamic size distribution either using fluorescence or scattering. The particles were diluted between $1 : 10^3 - 10^7$ in deinoized water such that the concentration during the measurement was close to $10^9$ particles per ml, where the deinoized water was first filtered through a 0.02 µm syringe filter (Whatman Anotrop, Cytiva) such that the particle count of the solution without particles was one or less per field of view. The sample was illuminated using a 488 nm laser module, and for the fluorescence measurements a 500 nm long-pass filter was inserted. All measurements were performed under flow to increase statistics and minimize bleaching, and at least 10 videos of each image modality were recorded.

The resulting size distributions can be seen in Figure S2. The NTA particle diameters were all within 20% of the size given by the manufacturer, where both the mode size and the standard deviation of the size distribution are similar to the size obtained from the CLiC measurements (see main text). For particles with diameter of 50 nm or larger, the scattering and fluorescence tracking have similar particle sizes. For measurements on samples with an average diameter below 50 nm, the resulting distributions from the fluorescence measurements are narrower and skewed toward smaller sizes compared to those obtained from the scattering signal. This indicates a missed subpopulation in the scattering channel as the particle sizes approach the detection limit.

#### 1.5 CLiC flow cell cleaning, surface treatment, and assembly

As previously reported[3, 4], CLiC uses a flow cell (Figure 1 in the main text and Figure S3) composed of two glass substrates (Ted Pella, product no. 260452, 200 ± 10µm thick, 25 mm × 25 mm cover glass) separated by a 30 µm thick double-sided adhesive (Nitto Denko, product no. 5603). The bottom glass substrate consists of an array of microfabricated cylindrical wells (0.5 µm deep × 3 µm diameter) designed for trapping samples (inset in Figure 1A (i - iii) in the main text), where the wells were fabricated by standard photolithography and dry etched by reactive ion etching (RIE)[5].

To prevent non-specific interactions for the reference dielectric bead measurements, the glass substrates were thoroughly cleaned using 1% Hellmanex solution in DI water followed by sonication in acetone and isopropanol, and a 45-minute piranha cleaning (a 3:1 mixture of sulfuric acid and hydrogen peroxide) just before flow cell assembly and the subsequent measurement. This cleaning approach creates a charge repulsion between the particle and the glass surfaces that is strong enough to prevent particle binding. For LNP measurements, the glass substrates were passivated shortly after piranha cleaning with 5 KDa PEG layers via silane chemistry and a cloud point PEGylation technique, previously reported[4, 6]. Briefly, the piranha-cleaned substrates were mildly-etched with a 1 M KOH solution and then exposed to vapor-phase deposition of 3-(aminopropyl)triethoxysilane (APTES) inside a vacuum desiccator at approximately 1.5 Torr for 30 minutes at room temperature. This step introduces primary amine groups onto the glass surface. A solution of sulfo-SMCC (4-(N-maleimidomethyl)cyclohexane-1-carboxylic acid 3-sulfo-N-hydroxysuccinimide ester) at 2 mg/mL in 1× diluted phosphate-buffered saline, PBS (15 mM salt concentration) is applied to the aminated substrates for 30 minutes at room temperature. This cross-linker reacts with the amine groups, introducing maleimide functionalities. The substrates were then immersed in a solution of thiol-terminated PEG (5 kDa) in

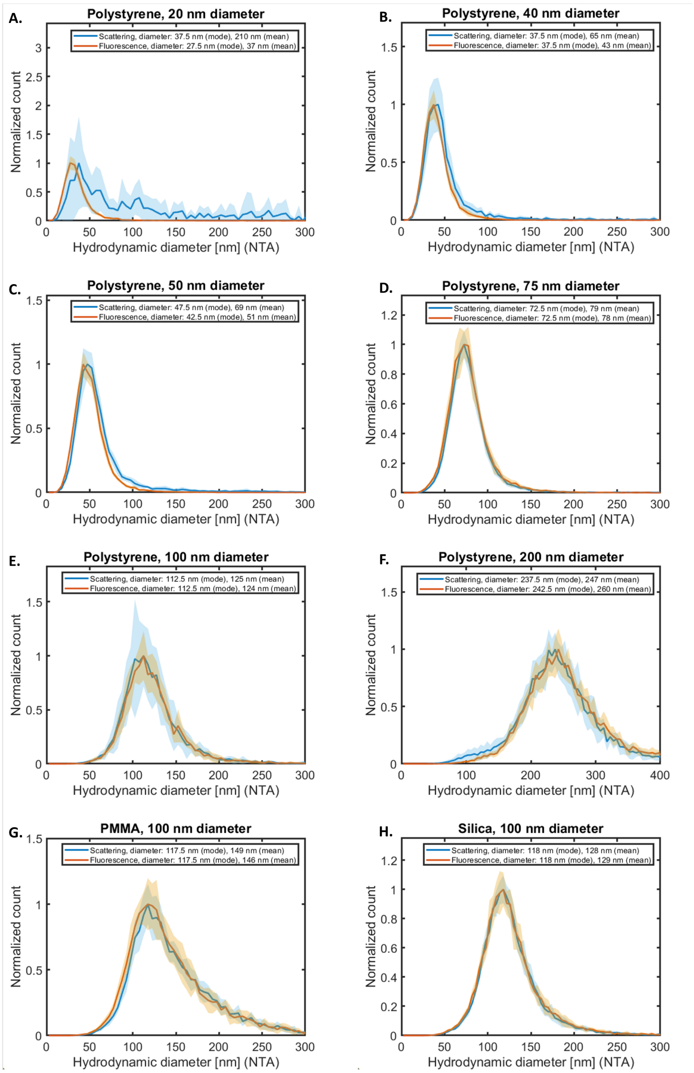

**Fig. S2 NTA size distribution of the used dielectric reference beads.** For each reference particle solution, the size distribution was measured both using the scattering signal and the fluorescence signal. The solid lines in the plots are the mean distribution for each modality (from at least 10 videos), where the shading is the standard deviation in each bin for all recording for each image modality. **A.** 20 nm diameter polystyrene, **B.** 40 nm diameter polystyrene, **C.** 50 nm diameter polystyrene, **D.** 75 nm diameter polystyrene, **E.** 100 nm diameter polystyrene, **F.** 200 nm diameter polystyrene, **G.** 100 nm diameter PMMA, and **H.** 100 nm diameter silica.

0.9 M $Na_2SO_4$ for at least 30-minutes. The use of $Na_2SO_4$ brings the solution close to the PEG's cloud point, a temperature at which PEG becomes less soluble, promoting a denser and more uniform PEG layer on the surface[6]. After PEGylation, the substrates were thoroughly rinsed with deionized water and dried with high-purity

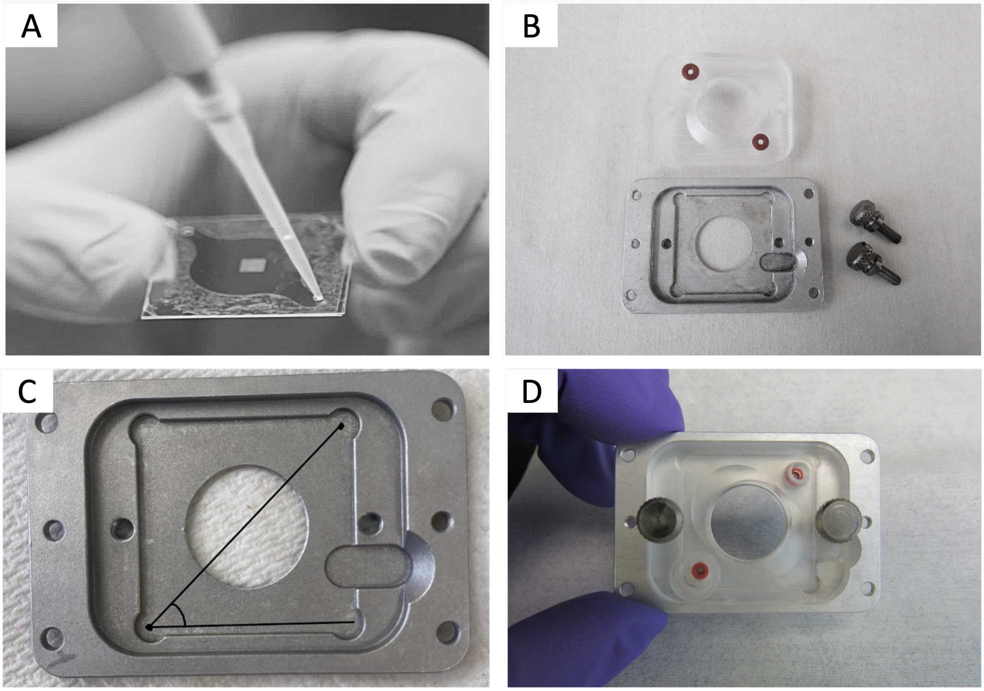

**Fig. S3 Images of the CLiC flow cell assembly and loading of the particle solution. A.** The assembled CLiC flow cell. The top coverslip has microfluidic inlets whereas the array of microwells is in the middle of the bottom coverslip. The two coverslips are sandwiched together by double-sided adhesive tape which forms the walls of the flow cell. **B-D** The baseplate and plastic chuck for connecting the assembled flow cell and tubing onto the microscope stage.

nitrogen before assembly. The glass coverslips were sandwiched together using the double-sided tape and assembled as shown in Figure S3.

##### 1.5.1 Choice of well diameter and depth

The used wells in the main text are cylindrical with a diameter of 3 μm and a depth of 0.5 μm. The choice of diameter comes from considering the number of wells in the field of view while also having large enough wells to be able to capture the particle diffusivity, where the latter is related to the slope in the MSD curve before reaching the plateau value (Figure 1 in the main text). Note that the diameter of the well used is the same as in previous work where the size of similar nanoparticles was accurately estimated using CLiC[4].

The choice to use 500 nm deep wells was based on two main considerations: 1) matching the focal depth of the confocal microscope to ensure particles remain in focus, and 2) minimizing the potential impact of confinement on diffusion. The former is particularly important when performing quantitative confocal microscopy, as the estimated particle signal otherwise would also depend on whether or not it is in focus. The latter is important because the depth of the well must be larger than the particle size so that the measured diffusivity can be related to the bulk diffusivity by using a correction factor based on the theory of particle diffusivity near planar surfaces. Using 500 nm wells ensures that the calibration factor is small, reducing the potential for systematic error. Using shallower wells, such as in the 100-200 nm regime, would restrict the ability to measure larger particles, as in Figure 4 in the main text. This need for calibration factors to relate confined diffusion to free-space diffusion is common to most methods that rely on particle confinement.

#### 1.6 Simultaneous CLiC-MC-iSCAT confocal Microscopy

The CLiC instrument (SCI100A-KIT Single-particle imaging technology package, ScopeSys) with an assembled microfluidic flow cell was placed on top of a microscope translation stage, of an inverted confocal microscope (Nikon AXR Ti2E), with the CLiC flow cell located between the imaging objective lens and a CLiC pusher lens, as illustrated in Figure 1 of the main text. Multichannel simultaneous fluorescence and label-free iSCAT experiments were performed utilizing the commercial Laser Scanning Confocal Microscope (Nikon, Japan) together with the integrated CLiC imaging platform, where the details of the CLiC instrument are previously described in[3, 7, 8]. We achieved both fluorescence and scattering detections by replacing the main dichroic mirror of the confocal microscope commonly used for separating the excitation and detection beam paths by a 20/80 (Reflected/Transmitted) beam splitter. The light from all imaging lasers are scanned simultaneously across the sample using the built-in resonant scanning mirrors, where the used microscope objective was a Nikon Plan 60x/1.42 NA oil immersion objective. During an experiment, the CLiC pusher lens was lowered to deflect and bring the flow cell surfaces into contact, hence trapping single diffusing particles into 3 µm diameter by 500 nm deep wells (within the focal plane of the imaging objective). To minimize the huge reflection signals from the top and bottom glass coverslips, we applied imaging oil in between the top glass coverslip and the CLiC pusher lens to create an index-matched optical path. For the iSCAT imaging, the high numerical aperture objective collected both the incident light reflected at the interface between the top and bottom glass coverslips and the imaging medium, as well as the backscattered light from the sample. The pinhole size was set to 2.0 A.U. to spatially filter the collected signals and detected by a photomultiplier tube (PMT). We installed a long-pass filter (T425LPXR, Chroma) and a clean-up filter (ZET405/20x, Chroma) in the filter cube of the turret sitting in front of the iSCAT PMT detector.

Moreover, an appropriate neutral density filter (ND Filter 10% Transmission, Chroma) was installed in the filter cube to reduce the amount of reflected light entering the iSCAT PMT. This modification allowed the incident 405 nm beam to lie within the bandpass of the filters in order to measure the intensity of the total light field. Similarly, fluorescence imaging was performed using any or all of the three laser wavelengths (488, 561, and 640 nm) along the same excitation path onto the three separate PMTs (emission filters 524/42 and 600/45 nm, Chroma) illustrated in Figure 1 in the main text. To minimize photobleaching of fluorescent dyes, experimental parameters included; exposure time of 20 ms and the maximum laser power of 0.5 mW, used in our measurements of the various LNPs formulations. With the separate detectors in place, hundreds to thousands of single particles are imaged and tracked simultaneously via fluorescent and label-free imaging, shown in Figure 2A in the main text for high throughput (Supplementary Movie 1–4). Each recorded video consists of 1000 frames, acquired over $\sim$ 20 seconds. The full field of view (FoV) captured during each scan is approximately 143 $\times$ 22 µm (1024 $\times$ 154 pixels at 0.14 µm/pixel) spanning about 24 individual microwells. Figure 2A and the supporting videos are cropped FoVs for best visualization. The tracked videos of confined fluorescently-labeled single-particle trajectories in time are extracted for quantitative measurements of diffusivity and size determination (Supplementary Movie 5).

During measurements, the particles were diluted within a range of $1 : 10^3 - 10^7$ to achieve a concentration where approximately 10-40% of wells contained an LNP. Reference particles were diluted in MilliQ water, while LNPs were diluted either in 1$\times$ PBS or 25 mM sodium acetate buffer. For CLiC measurements, following each recording, the microscope was programmed to automatically raise the CLiC lens and top coverslip to repopulate the particles and later released to bring the coverslips in contact again to trap fresh particles (Figure 1 in the main text). In most measurements, controlled dilution yields $\sim$ 5 to 15 particles in the field of view. Importantly, the platform is scalable in throughput: the experiment is repeated 100 to 200 times to increase the particle statistics and ensure robust data collection. The image analysis takes around 1 minute per video. This flexibility allows us to adapt the experiment for high-throughput screening when desired, without compromising single-particle resolution.

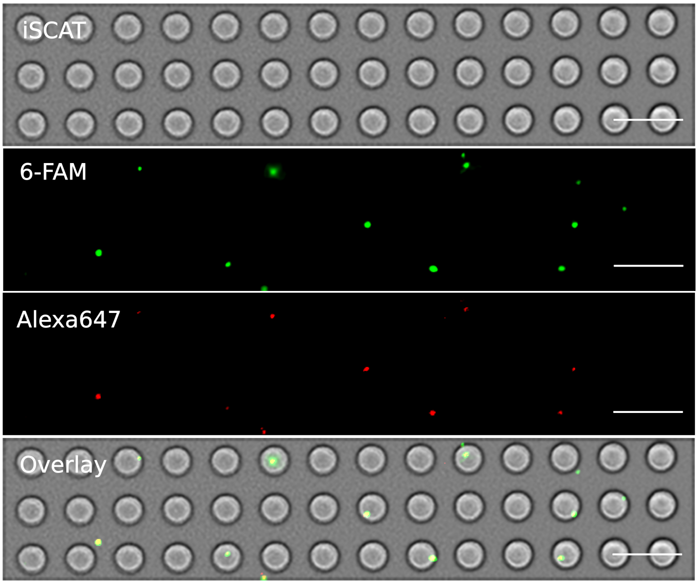

**Fig. S4 Simultaneous CLiC-MC-iSCAT imaging of lipid nanoparticles containing conjugated dual-labeled DNA complex.** Representative images from the iSCAT, 6-FAM and Alexa647 channels, respectively, showing dual fluorescence labeling of the DNA duplex at the formulating pH.7.4. Scale bar is 10 µm.

#### 1.7 CLiC image analysis

Data were analyzed using custom analysis software developed in MATLAB. The wells were identified by using edge detection on the raw iSCAT images, and subsequently a find circles algorithm was used. Only particle detections inside the wells were used for further analysis. To reduce the noise in the image, the images were convoluted with a Gaussian with a width of 1.5 pixels. Particle detection and in-plane subpixel localization of detected local extrema was performed using the radial center method[9] on the DiO signal for LNPs or the single-color fluorescence channel for the dielectric reference particles. Here the used signal threshold for particle detection is based on the standard deviation in the image. Maximum one particle detection per well was tracked, where in the case of more than one particle per well the brightest particle signal was tracked. However, the particle concentration was tuned such that it was mostly a single particle per well. Following particle detection in each frame, single-particle trajectories were derived by using a distance metric to associate detections from individual particles in previous frames. The observations were joined into traces by minimizing the sum of this metric using the Hungarian algorithm. For each included particle in the main text, the minimum track length was 120 frames.

Particle co-localization using other imaging channels was done by cropping a region of interest (ROI, $32 \times 32$ pixels) centered around the particle position in the fluorescence channel for all other imaging channels. For a completed particle trace, these ROIs were averaged and the particle signal was estimated by fitting the image to a 2D-Gaussian. Given that the particles were always in focus due to CLiC, such averaging where the particle was in the center of the ROI allowed to improve the signal-to-noise ratio of weak signals by effectively increasing the exposure time without inducing motion blur. For fluorescence co-localization, only the first 20 particle observations were averaged due to photobleaching. For iSCAT and DIC, all particle detections were used in the signal averaging, where the absolute values of ROIs were averaged because the depth motion of the particles influence both the amplitude and sign of the measured particle signal (Section 1.7.2).

To determine if a particle was co-localized with a second channel, the position-based averaged image in the second channel was fitted to a 2D-Gaussian. If the R-square value of the fit was larger than 0.5, the particle was considered co-localized. Moreover, for fluorescence, to minimize the risk that signal cross talk affects the estimation of the subpopulation containing mRNA, the Cy5-mRNA signal needed to be at least 2% of the DiO signals, where the threshold of 2% was estimated based on the used fluorophores, dichromatic mirrors, and emission filters.

##### 1.7.1 Handling of the iSCAT and DIC images

For the iSCAT and DIC images, a background buffer consisting of images before and after the image was used to subtract the background and normalize the signal in each well. Specifically, considering frame number $n$, the background buffer consists of $(n-5, ..., n-1, n+1, ..., n+5)$, where the frames just before or after the current frame are removed to avoid self-subtraction of the particle. The mean value of this background buffer is then used as the background frame. Thus, the contrast $C$ in the images after the background subtraction and normalization was defined

$$C = \frac{I_{\mathrm{n}} - I_{\mathrm{bg}}}{I_{\mathrm{bg}}}. \tag{1}$$

For iSCAT, to handle that the background signal may vary between wells due to differences in depth, which affect the background signal but not the number of photons scattered from the particle, the contrast was further multiplied with the factor $\sqrt{I_{\mathrm{bg}}/I_{\mathrm{const}}}$[10]. With this data handling, when the radius of the particle is much smaller than the wavelength of light, the contrast is proportional to the radius to the power of three[11]. To have a straightforward comparison between the particle signal and the particle radius, what was plotted in the figures are the cube root of the integrated scattering signal, where the integration was performed by fitting a Gaussian to the absolute value of the contrast image. The absolute value of the signal was used to take into account depth dependence of the iSCAT and DIC signal, where the signal can change sign as it diffuses in the depth direction, which limits the ability of averaging co-localized images to improve the signal-to-noise ratio. Thus, the estimated single-particle iSCAT contrast here represents the average absolute contrast value for all different depth positions the particle can have inside the well (Section 1.7.2).

##### 1.7.2 Depth position dependence and uncertainty in iSCAT signal estimation

The iSCAT signal is the interference between the particle signal and the local background. Due to the backscattering measurement geometry, the relative phase between the particle signal and the background here depends on the depth position of the particles[11]. This can, for example, be used to track the position of particles in 3D[12]. However, this requires that the particles are detectable in each frame, which is not the case here for all measured particles. Due to the comparatively weaker iSCAT signal compared to the fluorescence signal observed for the smaller nanoparticles discussed in the main text, particle tracking was performed using the fluorescence signal. The corresponding iSCAT signal was then estimated by position-based averaging of the absolute iSCAT contrast values, as described in (Section 1.7). As the absolute value of the iSCAT signal can be approximated as $|\sin(\frac{4\pi n_m \Delta z}{\lambda})|$, where $n_m$ is the refractive index of the medium and $\lambda$ is the wavelength in vacuum, it is expected that the average iSCAT signal is about 36% less than when using the maximum particle signal from a single frame.

Moreover, since the iSCAT signal depends on the depth position of the particles, it becomes important that the particles explore a wide range of different depth positions for the iSCAT signal to be accurately estimated. Since the iSCAT illumination wavelength is 405 nm in vacuum, the corresponding wavelength in water ($n \approx 1.34$) is approximately 300 nm. Given that the absolute iSCAT signal is periodic with a relative phase shift of $\pm\pi$, this corresponds to a z-movement of about 75 nm. Considering that the measured particles have diameters around 100 nm and the depth of the wells are 500 nm, the particles can move around 400 nm in axial position, which corresponds to about 5 periods of the iSCAT signal. Given that the minimum included track length is 120 particle observations, it can be assumed that the particle in the well explores all different depth positions it can be located at inside the well during the measurement. However, the depth-position range that the particle explores might not necessarily correspond to a complete period of the iSCAT signal. This in turn can cause a small bias in the iSCAT signal estimate based on the diameter of the particle. However, when calculating the estimated bias for the average iSCAT contrast from long particle track lengths, as long as the particles have at least 100 nm to move in depth, the potential bias is less than 6% (Figure S5). Thus, the spread in the estimated iSCAT signal in the main text originating from the particle estimate used corresponds to about 6%.

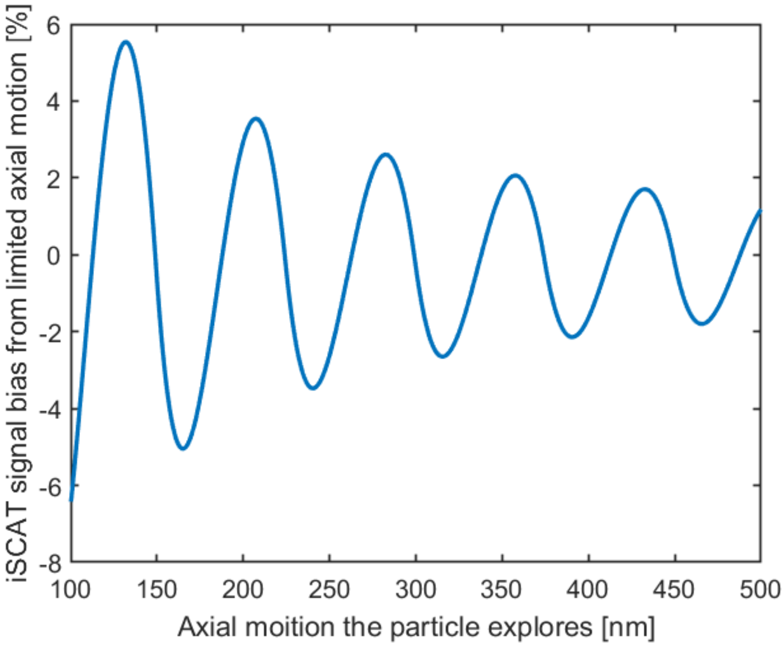

**Fig. S5 Potential iSCAT signal bias based on the limited axial movement.** Since the iSCAT signal depends on the particle's z-position, the depth-position range that the particle explore might not necessary correspond to a complete period of the iSCAT signal, there will be a small bias in the estimated iSCAT signal. However, for the expected movement ranges occurring in the main text, the bias is estimated to be less than 6%.

##### 1.7.3 Relating the fluorescence signal to particle properties

Since CLiC allows for fluorescence measurements without any pre-bleaching, it has been used for detailed investigations between the relation between fluorescence signal and size[4]. When using CLiC to measure colocalization of multiple fluorescence channels, such as DiO and Cy5-mRNA, the use of position-guided averaging minimizes the risk that signal contributions from potentially free mRNA affecting the loading estimate. This comes from the fact that only Cy5-mRNA that moves together with the LNP during the imaging will create a clear spot in the averaged image when the positional-guided averaging is based on the DiO signal. If there were some free mRNA in the wells together with an LNP, the free mRNA would mainly contribute to the average background signal in the averaged Cy5-mRNA image, where the background signal would not be picked up by the Gaussian fit to the image. Moreover, if the free mRNA would coincide with the LNP signal for some frames, there would be a continuum of Cy5 signals rather than the observed signal difference in the measured signal between loaded and empty LNPs.

Due to the lack of pre-bleaching, the Cy5-mRNA signal could in principle be used to measure the number of Cy5-mRNA copies inside the LNPs by also measuring the fluorescence signal from the signal from a single free mRNA. However, to convert the Cy5 signal to loading when using simultaneousness imaging, one needs to consider potential fluorescence proximity effects such as the FRET between DiO and Cy5 and fluorescence self-quenching. Thus, the precision of such copy number estimates depends on the available information of such fluorescence effects, which is why no copy number estimate is performed in this work.

#### 1.8 Theoretical size-iSCAT signal curve

A custom MATLAB script based on Mie calculations[13] was used to generate the theoretical size-iSCAT signal curve enabling estimation of the refractive index based on the measured iSCAT contrast and the particle size obtained from the single-particle tracking, where the used code is an adaptation of the freely available MatScat MATLAB package[13]. The calculations assume an angular distribution of the incoming light corresponding to the NA of the objective lens and that all light collected by the objective will reach the detector, where only the interferometric signal contribution is the considered. This approximation enables us to consider the two glass-fluid back reflections as a scaling of the background, which only affects the interferometric signal with a constant factor. The presence of two back reflections here increases the overall background as the two reflections are expected to interfere constructively based on the well depth and the wavelength used, which slightly decreases the relative contribution of the pure scattering term.

The incoming light at each angle is here approximated as a planewave with the amplitude of 1. For each such incoming planewave the corresponding Mie scattering[13] was calculated for the angles that can be captured by the objective based on the illumination. The scattering for all incoming angles is then summed and weighted based on the solid angle for each illumination angle. The parameters for the calculation were chosen to match those of the measurements. Specifically, the iSCAT wavelength (in vacuum) was 405 nm, the refractive index of the surrounding media (water) was assumed to be 1.34, and the captured angular range in water was assumed to be 85 degrees. To relate the Mie calculations to the measurements, there is only one free scaling factor which needs to be obtained from calibration measurements. The calibration of the Mie calculations to the experimental data was done using the 110 nm polystyrene sample by setting the mean estimated refractive index to 1.60.

In the case of widefield iSCAT, size vs iSCAT signal have local maxima and minima[10, 11, 14, 15], which is not present in the plots in the main text. This comes from the wide range of continuous distribution of illumination angles in the case of confocal iSCAT. Since the position of the local maxima and minima depends on the angle between the incoming and measured outgoing light[11], a wide range of scattering angles blurs the effect of any maxima/minima. Thus, for high-NA confocal microscopy measurements, the wide range of incoming and scattered light that reach the detector blur out the local minima in the reference curve.

Moreover, in the theoretical iSCAT curve, the slope decreases with increasing particle size. This behavior comes from the scaling of scattering intensity; in the Rayleigh regime, scattering scales with particle volume, whereas in geometrical optics– which is valid for particles much larger than the wavelength of light, scattering scales with the projected particle area[16]. In the intermediate regime, interference between various scattering elements within the particle, as descried by the optical form factor in Rayleigh-Debye-Gans theory, lowers the signal compared to Rayleigh scattering. Thus, as the particle size increases, the cube root of the interferometric backscattering signal will transition from being linear to sublinear, as seen for the theory lines in the figures of the main text.

##### 1.8.1 Single-particle refractive index estimate

The single-particle refractive index estimates in the main text were performed by combining the information from the single-particle hydrodynamic radius and the iSCAT contrast estimate. Specifically, for a known particle size, the iSCAT signal increases with the refractive index of the particle. Thus, for each measured particle, an iSCAT-refractive index curve is calculated using the same parameters as in Section 1.8 and the hydrodynamic radius of the measured particle. The intersection between the theoretical iSCAT-refractive index curve and the measured iSCAT signal is then used as the estimate of the single-particle refractive index.

When estimating single-particle refractive index, it is important to consider that an 10% uncertainty in the size estimate (Section 1.9) will cause $\sim 30\%$ uncertainty in the refractive index estimate as it relates to the particle volume estimate[17]. Thus, the spread in relative particle refractive index (difference from the refractive index of the medium) depends considerably on the accuracy of the size estimate. For this reason, the median particle refractive index is used when reporting the estimated particle refractive index.

##### 1.8.2 Limitations of the theoretical signal-size relation

When deriving the used relation between particle size and iSCAT signal it is assumed that the optical field of the background signal has a higher amplitude than the optical field from the particle scattering, which is a common approximation during iSCAT particle analysis[11]. This approximation allows the particle darkfield signal to be assumed to be negligible, which allows us to only consider the contribution of the interferometric signal. Although the obtained theoretical curve describes the experimental data well, there is an upper size limitation where this approximation no longer holds. In prior work, polystyrene particle sizes up to 200-300 nm in diameter has be used in iSCAT particle characterization in which only the interferometric contribution was considered[10, 15]. Given that the darkfield signal does not have the size z-position dependence as the interferometric particle signal, where the average iSCAT signal is $\sim 36\%$ lower than the maximum single-frame iSCAT signal (Section 1.7.2), the size range of when darkfield can be neglected is slightly reduced compared to traditional iSCAT image analysis. For particles above this upper signal limit of iSCAT, transmission methods can still be accurately used due to the difference in signal-size scaling compared to backscattering methods[10], where it is for example possible to estimate size and refractive index directly from a transmission microscopy image for particles larger than 300 nm in diameter[18]. Given that particles with an optical signal corresponding to 100 nm diameter polystyrene or higher can be measured using DIC, this shows that the combined iSCAT and transmission microscopy has a larger dynamical range than iSCAT measurements alone[19]. Thus, for particles with an optical signal that is intense enough so that the traditional iSCAT approximation (that the optical field of the background signal has an higher amplitude than the optical field from the particle scattering) no longer holds, the compatibility with transmission methods for CLiC-MC-iSCAT becomes critical.

For the refractive index estimations of the LNPs, it is important that the limitation of neglecting the darkfield contribution is related to the scattering amplitude of the particles. Given that the scattering signal amplitude of the LNPs is less than or similar to that of the 100 nm polystyrene beads, where the obtained iSCAT-size relation seems to describe well both the 100 and 200 nm polystyrene beads, the approximation seems to be valid for the measured LNPs.

#### 1.9 Estimating particle size from trajectories

When relating the particle trajectories to hydrodynamic diameter, both the finite size of the well and the proximity between the particle and the surfaces in the well need to be considered. Details of how this was performed for the used geometry can be found in Ref.[4]. Briefly, each particle trajectory is first converted to a mean squared displacement (MSD) curve, and only particles with at least 120 detections are included in the analysis. The resulting MSD curve is then fitted using a model that describes diffusion in a circular well[4], with an example shown in Figure 1C of the main text. The mean squared displacement of a particle confined by a circle is described by[4]

$$\mathrm{MSD}(t) = O + r^2\Big(1 - 8\sum_{m=1}^{\infty} \frac{\exp\left(-\alpha_{1m}^2 t/\tau\right)}{\alpha_{1m}^2(\alpha_{1m}^2 - 1)}\Big), \tag{2}$$

where $O$ is a offset from position localization uncertainty, $r$ is the confinement radius, $\tau = r^2/D$ is the characteristic time, $D$ is particle diffusivity, and $\alpha_{1m}$ is the $m^{\mathrm{th}}$ positive root of the derivative of the Bessel function of the first kind. To estimate the diffusivity in this work, the first two terms in the expansion are used, which well describes the MSD obtained, as exemplified in Figure 1C in the main text. From that fit, the 2D diffusivity of the particle inside the well is extracted. However, due to confinement and the proximity to nearby surfaces, the diffusivity of the particle in the well is lower than the corresponding bulk diffusion constant[20].

To convert diffusivity to size distributions, we use a modified Stokes–Einstein relation[21]:

$$D_{||} = \frac{k_\mathrm{B}T}{6\pi\eta\lambda a} = \lambda^{-1}D_0, \tag{3}$$

where $D_{||}$ is the diffusivity of particles near two parallel confining planes, $D_0$ is the diffusivity in bulk, $k_\mathrm{B}$ is the Boltzmann constant, $T$ is the temperature, $\eta$ is the kinetic viscosity of the solution, and $a$ is the hydrodynamic radius. $\lambda$ is a correction factor used to account for the hydrodynamic effects near surfaces and can be approximated as[21, 22]

$$\lambda^{-1} = 1 - 1.004\Big(\frac{a}{z}\Big) + 0.418\Big(\frac{a}{z}\Big)^3 + 0.21\Big(\frac{a}{z}\Big)^4 - 0.169\Big(\frac{a}{z}\Big)^5 + O\Big(\frac{a}{z}\Big)^6, \tag{4}$$

where $z$ is the midway distance between the two planes of confinement. Given the known well depths (500 nm), the confinement effect is corrected by determining the correction factor that is self-consistent with the hydrodynamic diameter estimated from the Stokes-Einstein relation.

Note that the closer the particle is to a surface, the slower the diffusivity of the particle. Thus, if particle-surface of the well interaction would occur, the size of the particles would be consistently overestimated. Since the obtained particle sizes agree well with that of complementary darkfield NTA measurements, this further indicates that the used model well describes the motion of the particles inside the wells.

Regarding the potential effect of motion blur, although the exposure time per frame was 20 ms, due to scanning of the microscope, the local exposure time is much less than 20 ms. To evaluate potential motion blur in our imaging system, we estimated the 1D root-mean-square displacement of freely diffusing 100 nm polystyrene beads with a diffusion coefficient $\sim 5.0$ µm/s. The frame rate of our confocal resonant scanning system is 50 Hz, which corresponds to the frame time of 20 ms. Each frame is composed of 154 rows (Y), meaning that the time to scan a single row is $\sim 130$ µs. Within each row, the resonant scanner samples 1024 pixels in X, giving a pixel dwell time of $\sim 127$ ns. Our measured lateral point spread function (PSF) $\sim 230$ nm FWHM, which corresponds to $\sim 2$ pixels at our system calibration of 0.14 µm/pixel. Therefore, during a single row scan, the expected 100 nm particle motion is $\sim 36$ nm. This shows that motion blur per row is minimal (less than 0.3 pixels), hence does not significantly affect our spatial resolution or particle localization.

##### 1.9.1 Statistical uncertainty on the particle size estimate using single-particle tracking

Estimation of the hydrodynamic radius using single-particle tracking is based on estimating the diffusivity of the measured particle using the particle's mean squared displacement, where the diffusivity can in turn be related to particle size. Since the observed diffusivity of a particle is a stochastic process, the uncertainty of the diffusivity estimate depends on the track length of the particle. The relative uncertainty in the diffusivity estimate is $\approx \sqrt{1/N}$, where $N$ is the number of particle observations[23].

Given that the minimum track length for the included particles is here 120 observations, the statistical uncertainty in the particle size estimate for the particles in the main text is less than 10%.

#### 1.10 Interpretation of the water fraction estimates

The term "water content" in the manuscript includes both the water inside the LNP core and the water associated with the external PEG layer, as the water associated with the external PEG layer contributes to the hydrodynamic radius. Thus, the estimate includes both the internal solvent fraction and the hydration shell water, which makes it different from other estimates using, for example, SAXS and SANS, which typically only refer to the internal solvent fraction in the core of LNPs[24].

To understand the difference in the estimated water fraction when including the hydration layer, the outer surface of the LNPs contains a PEG layer, where a 2k PEG layer has a thickness around 4 nm[24]. For a particle with a diameter of 90 nm, the 4 nm PEG layer corresponds to 25% of the estimated particle volume. Because the PEG layer has a low bimolecular density, this indicates that around half of the estimated LNP water fraction in the main text corresponds to the water in the hydration shell.

Regarding the estimate of the water fraction of the LNPs that contain mRNA cargo, according to cryoTEM data, the measured LNPs contain so-called "blebs" (Figure S1). Blebs are associated with a water phase inside the LNPs, while LNPs without mRNA generally do not contain blebs[25]. Thus, the observation that the LNPs containing cargo have a lower refractive index than those without cargo is likely due to the fact that the water in the bleb will counteract the increased refractive index of the mRNA.

Regarding the increased estimated water fraction of the LNPs at reduced pH, in prior work[26] it has been hypothesized that electrostatic repulsion between charged MC3 lipids at reduced pH triggers a structural change in which the resulting influx of water and ions balances the repulsion. This comes from that the N/P ratio here is 3, which means that charges from the MC3 lipids will be more than those from the mRNA. Due to the net positive charge inside the LNPs, there will be an electrostatic repulsion between the MC3 lipids, which can be counteracted by an influx of water and ions.

#### 1.11 Comparison with other particle sizing techniques

A growing number of analytical techniques have emerged to characterize the size and composition of nanoparticles at the single-particle level. Our CLiC-MC-iSCAT offers a powerful approach for real-time tracking of freely diffusing particles under confined conditions, enabling simultaneous label-free and fluorescence detections for size estimation, refractive index, and fluorescence-based content quantification. This method complements a broad array of established and emerging nanoparticle characterization platforms including: interferometric nanoparticle tracking analysis (iNTA)[14], single-particle interferometric reflectance imaging sensor (SP-IRIS)[27], nanoparticle tracking analysis (NTA)[28], tunable resistive pulse sensing (TRPS)[29], nano flow cytometry (nFCM)[30], atomic force microscopy (AFM), cryogenic transmission electron microscopy (Cryo-TEM), and waveguide microscopy[17]. At the ensemble level, optical and scattering-based techniques like dynamic light scattering (DLS)[31] remain widely used for size estimation in solution. Table 1 provides a comparison of these methods, outlining their core principles, compatibility with solution-based measurements, and their ability to provide fluorescence and label-free detection, payload quantification, and extended observation of individual nanoparticles. In that comparison, it can be noted that although similar particle information can be obtained using other techniques, CLiC-MC-iSCAT obtains a wide range of particle information of particles in suspension without the need of a specialized optical setup, which makes the method different compared to the other methods.

**Table 1** Comparison of different particle size characterization techniques. Working principles of DLS, NTA, SP-IRIS, TRPS, AFM, Cryo-TEM, nFCM, iNTA, and CLiC-MC-iSCAT methods. CLiC is unique in enabling untethered single-particle tracking in solution with extended observation times of the same confined particles.

| **Technique** | **Solution** | **Principle** | **Special setup** | **Fluorescence** | **Size Payload** | **Prolonged Watch** |
|---|---|---|---|---|---|---|
| CLiC-MC-iSCAT | Yes | iSCAT and multi-color confined single particle tracking. | No | Yes | Yes | Yes |
| iNTA | Yes | iSCAT, tracking of diffusing particles. | Yes | No | Size | No |
| nFCM | Yes | scattering, fluorescence in flow-based measurements. | Yes | Yes | Yes | No |
| SP-IRIS | No | Interferometric imaging to detect and characterize immobilized nanoparticles. | Yes | Yes | Yes | No |
| NTA | Yes | tracking of Brownian diffusing particles. | Yes | Yes | Yes | No |
| DLS | Yes | time-dependent intensity fluctuations of diffusing nanoparticles. | Yes | No | Size | No |
| TRPS | Yes | measure changes in electrical resistance as single particles pass through a pore. | Yes | Yes | Size | No |
| AFM | No | topographic image of bound particles. | Yes | Yes | Size | Yes |
| Cryo-TEM | No | Electron scattering | Yes | Yes | Yes | No |
| Waveguide microscopy | No | Evanescent field scattering and fluorescence. | Yes | Yes | Yes | Yes |

### 2 Complementary data

#### 2.1 CLiC-MC-iSCAT size distribution of the measured reference beads

When performing CLiC-MC-iSCAT, particles can be tracked as long as at least one of the detection channels provides sufficiently strong signal. In the current setup, particles below a certain size are no longer detectable in iSCAT but remain detectable in fluorescence, which allows for sizing comparable to other single-particle tracking methods (Figure S6). For polystyrene particles, the iSCAT detection limit lies between 40 and 50 nm in particle diameter. Importantly, particle colocalization with other fluorescent labels, such as Cy5-mRNA, remains valid even when the iSCAT signal is not detectable, allowing continued investigation of particle-associated fluorescence properties.

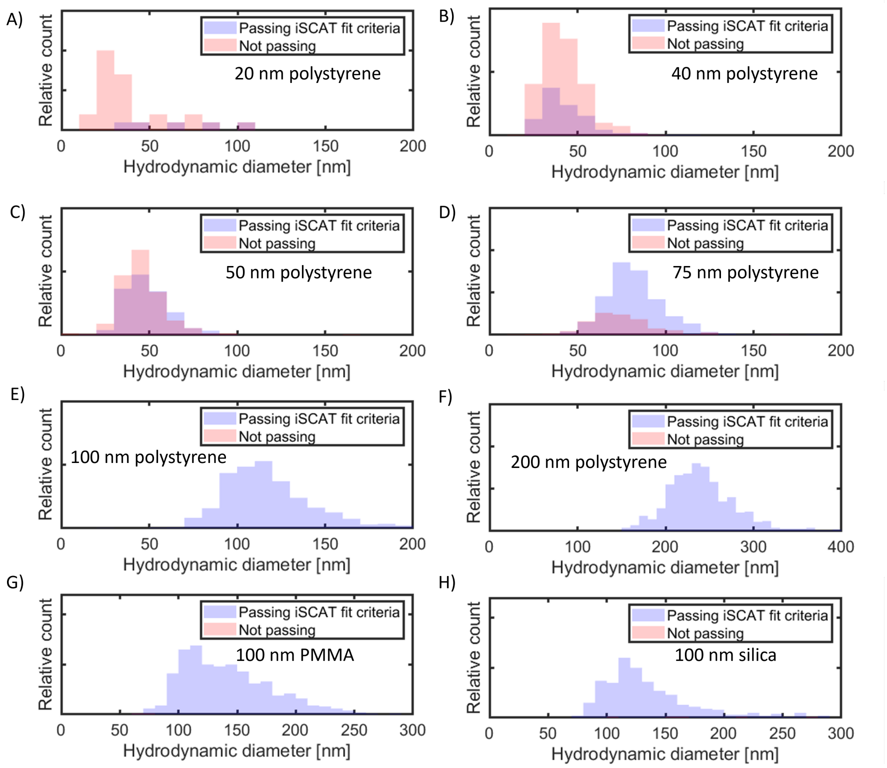

**Fig. S6 CLiC-MC-iSCAT size distribution of the used dielectric reference beads.** For each reference particle solution, the detected particles are divided into two different categories: those that have a detectable colocalized iSCAT signal and those that do not have that, but are still detectable in the fluorescence channel. **A.** 20 nm diameter polystyrene, **B.** 40 nm diameter polystyrene, **C.** 50 nm diameter polystyrene, **D.** 75 nm diameter polystyrene, **E.** 100 nm diameter polystyrene, **F.** 200 nm diameter polystyrene, **G.** 100 nm diameter PMMA, and **H.** 100 nm diameter silica.

#### 2.2 Evaluation of CLiC-MC-iSCAT using multi-labeled reference beads

To evaluate particle colocalization across multiple imaging channels, reference nanobead samples in which each particle is labeled with a set of four different fluorophores were measured with CLiC-MC-iSCAT (Figure S7). In that measurement, all detected particles are colocalized with a fluorescence signal in the other channels, as indicated by that the fluorescence signal ratio between the channels are close to one for all detected particles (Figure S7B). Moreover, the amplitude of the single-particle fluorescence all scale with the iSCAT signal. Combined, this indicates that when a particle contains several different fluorophores, CLiC-MC-iSCAT can accurately colocalize the different fluorescence signals and that the fluorescence signal reflects the amount of fluorophores per particle.

#### 2.3 Effect on measured size and iSCAT signal from potential particle swelling

To investigate the physical meaning of the change in the measured CLiC particle size and the iSCAT signal for DNA-containing LNPs at different pH conditions of the medium, the size of the LNPs was also measured using DLS (Figure S8). The measured size distributions using DLS and CLiC at the two different pH are similar, indicating that the increase in particle size is likely physical and not a potential artifact from a pH-dependent LNP-surface interaction during the CLiC measurement.

Furthermore, in Figure 4 in the main text, the iSCAT signal appears to be slightly reduced after swelling. To investigate whether this is an indication of mass loss or a consequence of the swelling, Mie calculations were performed where the particle refractive index was assumed to scale linearly with the bimolecular concentration inside the particle (Figure S9). Thus, changes in particle volume cause a corresponding change in particle refractive index. When simulating a particle with a refractive index of 1.44 when the diameter is 100 nm, the iSCAT signal reduces as the particle size increases. This is in line with the size-iSCAT signal plots in the main text where the slope decreases with increasing particle size. Thus, the observations of increased particle size and a slightly reduced iSCAT signal in Figure 4 in the main text are indications of particle swelling.

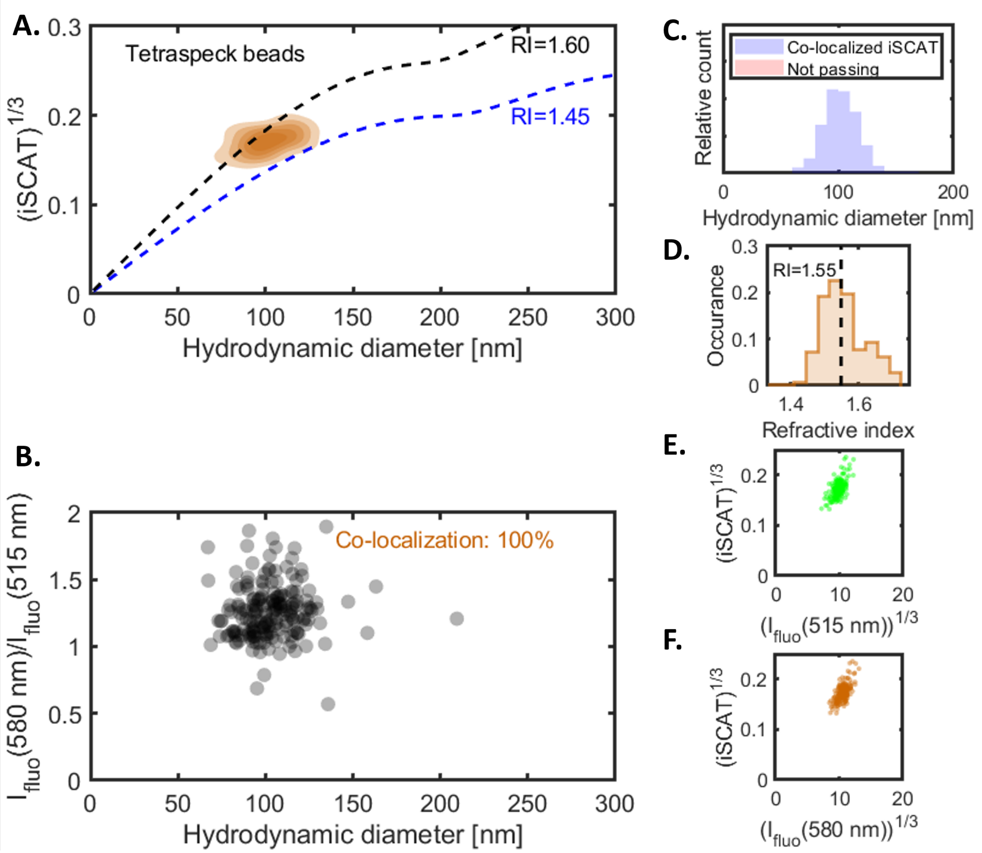

**Fig. S7 CLiC-MC-iSCAT measurement of multi-labeled reference beads.** The reference beads are labeled with four different fluorophores (100 nm diameter Tetraspeck beads), allowing for evaluation of the colocalization of multiple fluorescent signals. **A.** Size-iSCAT contour plot, where the signal is close to that of polystyrene. The contour lines correspond to 17, 33, 50, 67, and 83% of the obtained distribution density. **B.** Ratio between the 580 nm and 515 nm fluorescence emission channels. That all the ratios are within 0.5-2.0 indicate that all particles are accurately co-localized with multiple fluorescence signals. **C.** Size histogram, where the particles have a size around 100 nm diameter, which also is the size given by the manufacturer. **D.** Refractive index histogram, where the median refractive index of 1.55. **E.** iSCAT as a function of 515 nm fluorescence signal, where the two signals scale with each other as expected. **F.** iSCAT as a function of 515 nm fluorescence signal, where the two signals scale with each other as expected.

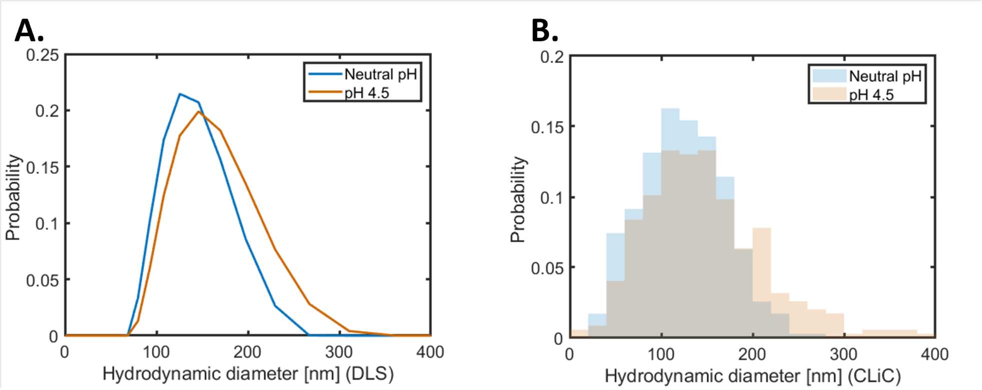

**Fig. S8 DLS and CLiC size distribution of LNPs at different pH.** The LNP size disquisition at neutral and reduced pH measured using **A.** DLS number average and **B.** CLiC. The size increase at the lowered pH compared to at neutral pH is similar for both techniques, indicating that the measured size increase using CLIC is physical and not an potential artifact from changed surface interaction.

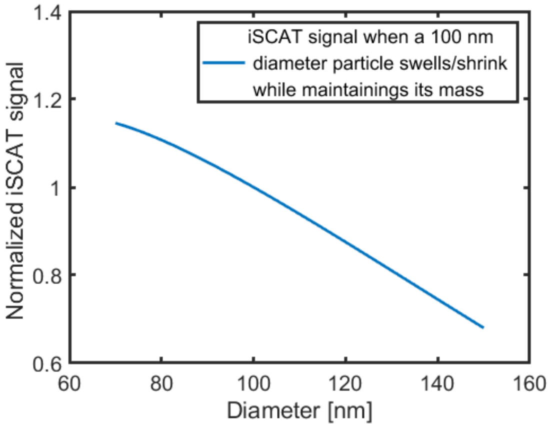

**Fig. S9 Theoretical change in measured iSCAT signal during particle swelling with maintained particle mass.** For these calculations, the particle refractive index was assumed to scale linear with bimolecular concentration. Thus, changes in particle volume cause a corresponding change in particle refractive index. Here, the particle has a refractive index of 1.44 when the diameter is 100 nm to mimic the LNP data. As an effect of swelling the iSCAT signal slightly drops, which is inline with the observation in Figure 4 in the main text.

[22] J. Happel and H. Brenner, *Low Reynolds number hydrodynamics: with special applications to particulate media*, vol. 1. Springer Science & Business Media,

2012.

[23] C. L. Vestergaard, P. C. Blainey, and H. Flyvbjerg, "Optimal estimation of diffusion coefficients from single-particle trajectories," *Physical Review E*, vol. 89, no. 2, p. 022726, 2014.

[24] M. Yanez Arteta, T. Kjellman, S. Bartesaghi, S. Wallin, X. Wu, A. J. Kvist, A. Dabkowska, N. Székely, A. Radulescu, J. Bergenholtz, *et al.*, "Successful reprogramming of cellular protein production through mrna delivered by functionalized lipid nanoparticles," *Proceedings of the National Academy of Sciences*, vol. 115, no. 15, pp. E3351–E3360, 2018.

[25] J. B. Simonsen, "A perspective on bleb and empty lnp structures," *Journal of Controlled Release*, vol. 373, pp. 952–961, 2024.

[26] Z. Li, J. Carter, L. Santos, C. Webster, C. F. van der Walle, P. Li, S. E. Rogers, and J. R. Lu, "Acidification-induced structure evolution of lipid nanoparticles correlates with their in vitro gene transfections," *ACS nano*, vol. 17, no. 2, pp. 979–990, 2023.

[27] F. Deng, A. Ratri, C. Deighan, G. Daaboul, P. C. Geiger, and L. K. Christenson, "Single-particle interferometric reflectance imaging characterization of individual extracellular vesicles and population dynamics," *Journal of Visualized Experiments*, no. 179, 2022.

[28] E. van der Pol, F. A. Coumans, A. Sturk, R. Nieuwland, and T. G. van Leeuwen, "Refractive index determination of nanoparticles in suspension using nanoparticle tracking analysis," *Nano letters*, vol. 14, no. 11, pp. 6195–6201, 2014.

[29] G. R. Willmott, "Tunable resistive pulse sensing: Better size and charge measurements for submicrometer colloids," *Analytical Chemistry*, vol. 90, no. 5, p. 2987–2995, 2018.

[30] S. Zhu, L. Ma, S. Wang, C. Chen, W. Zhang, L. Yang, W. Hang, J. P. Nolan, L. Wu, and X. Yan, "Light-scattering detection below the level of single fluorescent molecules for high-resolution characterization of functional nanoparticles," *ACS Nano*, vol. 8, no. 10, p. 10998–11006, 2014.

[31] H. C. V. D. Hulst, *Light Scattering by Small Particles.* Mineola, New York: Dover Publications Inc., 1981.
